# alignparse: A Python package for parsing complex features from high-throughput long-read sequencing

**DOI:** 10.1101/850404

**Authors:** Katharine H.D. Crawford, Jesse D. Bloom

## Abstract

Advances in sequencing technology have made it possible to generate large numbers of long, high-accuracy sequencing reads. For instance, the new PacBio Sequel platform can generate hundreds of thousands of high-quality circular consensus sequences in a single run (Rhoads and F. 2015; Hebert et al. 2018). Good programs exist for aligning these reads for genome assembly (Chaisson and Tesler 2012; Li 2018). However, these long reads can also be used for other purposes, such as sequencing PCR amplicons that contain various features of interest. For instance, PacBio circular consensus sequences have been used to identify the mutations in influenza viruses in single cells (Russell et al. 2019), or to link barcodes to gene mutants in deep mutational scanning (Matreyek et al. 2018). For such applications, the alignment of the sequences to the targets may be fairly trivial, but it is not trivial to then parse specific features of interest (such as mutations, unique molecular identifiers, cell barcodes, and flanking sequences) from these alignments.

Here we describe alignparse, a Python package for parsing complex sets of features from long sequences that map to known targets. Specifically, it allows the user to provide complex target sequences in Genbank format that contain an arbitrary number of user-defined sub-sequence features. It then aligns the sequencing reads to these targets and filters alignments based on whether the user-specified features are present with the desired identities (which can be set to different thresholds for different features). Finally, it parses out the sequences, mutations, and/or accuracy of these features as specified by the user. The flexibility of this package therefore fulfills the need for a tool to extract and analyze complex sets of features in large numbers of long sequencing reads.

## Uses & Examples

Below are two example use cases of alignparse from our research:

### Sequencing deep mutational scanning libraries

In deep mutational scanning experiments, researchers use mutant libraries to assay the effects of tens of thousands of individual mutations to a gene-of-interest in one experiment (Fowler and Fields 2014). One way to make deep mutational scanning of long gene variants work efficiently with short-read Illumina sequencing is to link the mutations in each variant to a unique molecular barcode (Hiatt et al. 2010). This barcode linking can be done by long-read PacBio sequencing of the variant library (Matreyek et al. 2018), but it is then necessary to parse the resulting long reads to associate the barcode with the mutations in the variant.

The alignparse package provides a standard tool for parsing barcodes and linked mutations from the long-read sequencing data. It also allows for the parsing of additional sequence features necessary for validating the quality of deep mutational scanning libraries, such as the presence of terminal sequences or other identifying tags. The RecA deep mutational scanning library example demonstrates this use.

### Single-cell viral sequencing

Some viral genomes are sufficiently small to be sequenced in their entirety using long-read sequencing technology. Recent work has shown that such long-read sequencing of viral genes can be combined with standard single-cell transcriptomic technologies (such as 10x Chromium) to simultaneously sequence the infecting virus and characterize the transcriptome in single infected cells (Russell et al. 2019). Such experiments require parsing the long-read viral sequences to identify viral mutations as well as cell barcodes, unique molecular identifiers, and other flanking sequences. The single-cell virus sequencing example shows how such parsing can readily be performed using alignparse.

## How alignparse works

alignparse takes the following inputs:

1. A user-defined Genbank file containing the sequence of one or more alignment targets with an arbitrary number of user-defined features.
2. A YAML file containing parsing specifications for each feature. These specifications include filters indicating the maximal allowed mutations in each feature, as well as information on what output should be parsed for each feature (e.g., its sequence, its mutations, or simply if it is present)
3. A FASTQ file containing the long-read sequencing data.

These inputs are used to define a Targets object. alignparse then uses this Targets object to create sequence alignments and parse sequence features defined in the input Genbank and YAML files.

alignparse aligns sequencing reads to the targets using minimap2. The alignparse.minimap2 submodule provides alignment specifications optimized for the two example use cases described above. alignparse uses the cs tags generated by minimap2 to extract the relevant features from the alignments into intuitive data frames or CSV files.

Downstream analyses of parsed features are facilitated by the alignparse.consensus submodule. This submodule provides tools for grouping reads by shared barcodes, determining consensus sequences for barcoded reads, and further processing mutation information for downstream analyses. Since the main outputs from alignparse are in intuitive data frame formats, downstream analyses can be highly customized by the user. Thus, alignparse provides a flexible and useful tool for parsing complex sets of features from high-throughput long-read sequencing of pre-defined targets.

## Code Availability

The alignparse source code is on GitHub at https://github.com/jbloomlab/alignparse and the documentation is at https://jbloomlab.github.io/alignparse.

## Acknowledgements

We would like to thank members of the Bloom lab for helpful discussions and beta testing. This work was supported by the following grants from NIAID of the NIH: R01 AI141707 and R01 AI140891. JDB is an Investigator of the Howard Hughes Medical Institute.

## References

Chaisson, M. J., and G. Tesler. 2012. “Mapping Single Molecule Sequencing Reads Using Basic Local Alignment with Successive Refinement (Blasr): Application and Thory.” BMC Bioinformatics 13 (238). https://doi.org/10.1186/1471-2105-13-238.

Fowler, D. M., and S. Fields. 2014. “Deep Mutational Scanning: A New Style Ofo Protein Science.” Nature Methods 11 (8). https://doi.org/10.1038/nmeth.3027.

Hebert, P. D. N., T. W. A. Braukmann, S. W. J. Prosser, S. Ratnasingham, J. R. deWaard, N. V. Ivanova, D. H. Janzen, et al. 2018. “A Sequel to Sanger: Amplicon Sequencing That Scales.” BMC Genomics 19 (219). https://doi.org/10.1186/s12864-018-4611-3.

Hiatt, J. B., R. P. Patwardhan, E. H. Turner, C. Lee, and J. Shendure. 2010. “Parallel, Tag-Directed Assembly of Locally Derived Short Sequence Reads.” Nature Methods 7 (2). https://doi.org/10.1038/nmeth.1416.

Li, H. 2018. “Minimap2: Pairwise Alignment for Nucleotide Sequences.” Bioinformatics 34 (18): 3094–3100. https://doi.org/10.1093/bioinformatics/bty191.

Matreyek, K. A., L. M. Starita, J. J. Stephany, B. Martin, M. A. Chiasson, V. E. Gray, M. Kircher, et al. 2018. “Multiplex Assessment of Protein Variant Abundance by Massively Parallel Sequencing.” Nature Genetics 50: 874–82. https://doi.org/10.1038/s41588-018-0122-z.

Rhoads, A., and Au K. F. 2015. “PacBio Sequencing and Its Applications.” Genomics, Proteomics & Bioinformatics 13 (5): 278–79. https://doi.org/10.1016/j.gpb.2015.08.002.

Russell, A. B., E. Elshina, J. R. Kowalsky, A. J. W. te Velthuis, and J. D. Bloom. 2019. “Single-Cell Virus Sequencing of Influenza Infections That Trigger Innate Immunity.” Journal of Virology 93 (14): e00500–19. https://doi.org/10.1128/JVI.00500-19.

